# Microbial metabolites mediate bumble bee attraction and feeding

**DOI:** 10.1101/549279

**Authors:** Robert N. Schaeffer, Caitlin C. Rering, Isabelle Maalouf, John J. Beck, Rachel L. Vannette

**Author notes:** Corresponding author and current address: Department of Biology, Utah State University, Logan, UT 84321. Equally-contributing authors (Robert N. Schaeffer, Caitlin C. Rering).

## Abstract

Animals such as bumble bees use chemosensory cues to localize and evaluate essential resources. Increasingly, it is recognized that microbes can alter the quality of foraged resources and produce metabolites that act as foraging cues. The distinct nature of these sensory cues however and their use in animal foraging remain poorly understood. Here, we test the hypothesis that species of nectar-inhabiting microbes differentially influence pollinator attraction and feeding via microbial metabolites in nectar. We examined electrophysiological potential of bumble bee antennae to respond to volatile microbial metabolites, followed by behavioral responses using choice assays. We assessed gustatory responses through both no-choice and choice feeding assays. Antennae responded to some microbial volatiles, and bees chose *Asaia* bacterial solutions compared to *Metschnikowia* yeast based on volatiles alone. However, *B. impatiens* consumed significantly more Metschnikowia-inoculated nectar, suggesting distinct roles for volatile and non-volatile microbial metabolites in mediating feeding decisions, with potential to affect associative learning and future foraging. Our results suggest that microbial metabolites may represent non-reinforcing cues with potential consequences for forager learning, economics and floral host reproduction.

## Introduction

To successfully persist in a chemosensory environment, animals must receive and interpret cues and signals of ecologically-important information, such as the quantity and quality of resources potentially available to them [1]. This is especially true of pollinators such as bumble bees, which integrate multi-modal signals, including form, color, and scent, to accurately identify rewarding flowers [2]. Like other food resources, flowers host varied microbial species and communities [3,4], which produce metabolites that may act as cues of resource availability and quality, with consequences for pollinator foraging [5,6]. Indeed, insect pollinators are highly sensitive to shifts in volatile abundance and identity [7–9], with scents being known to influence learned foraging preferences [10]. However, the role of volatile and non-volatile microbial metabolites in mediating pollinator attraction and foraging decisions still remains largely unclear.

In standing crop nectar, bacteria and fungi colonize between 20-70% of individual flowers, attain densities exceeding 10^5^ and 10^7^ cells/μL respectively [3,4] and metabolize sugars and amino acids [5,11], affecting pollinator foraging and plant reproduction [5,12,13]. Intense competition between microbes in nectar often results in flowers that are dominated by either yeast or bacteria [14]. Yeasts and bacteria differ in volatile composition and acceptance to pollinators [6], but also differentially influence non-volatile nectar traits (Vannette & Fukami 2018) and shift pollinator perceptions of nectar quality [15]. Predicting microbial effects on pollinator foraging and behavior requires examining responses to olfactory (headspace volatiles) and gustatory (dissolved chemicals) cues.

Here, we test the hypothesis that yeast and bacteria differentially influence bumblebee attraction and feeding. Bumble bees (*Bombus impatiens*) are an ideal system, due to their close ecological relationships with yeasts [16,17] and bacteria [18,19]. We examined antennal responses to microbial metabolites using electroanntenographic (EAG) bioassays, bee choice using olfactometer (Y-tube) bioassays, and gustatory preferences using choice and no-choice feeding assays. We found bumble bees show distinct responses to volatile vs gustatory microbial cues to inform foraging decisions, indicating the potential for associative learning, where bumble bees may adjust behavioral responses to volatile blends after exposure to gustatory microbial cues.

## Materials and methods

### Study system

We used three colonies of the generalist bumble bee *Bombus impatiens* (Koppert Biological Systems, Inc.; Howell, MI, USA) and strains of the nectar-inhabiting yeast *Metschnikowia reukaufii* (Metschnikowiaceae; GenBank ID: MF319536) and bacteria *Asaia astilbes* (Acetobacteraceae; GenBank ID: KC677740). Both *M. reukaufii*, and *A. astibles* are commonly isolated from floral nectar [20] and pollinators (Good *et al*. 2014), but differentially influence nectar chemistry and scent [21].

### Experiment 1: Electroantennographic bioassay

We examined antennal response (n=6 /metabolite) to volatiles produced by *Metschnikowia* and *Asaia* (Table 1) by puffing each metabolite (0.4 μmol) over excised *B. impatiens* antennae. Antennal responses were recorded and corrected by responses to blanks and positive control stimuli (0.4 μmol geraniol), see electronic supplementary material S1 Methods.

**Table 1 -.**
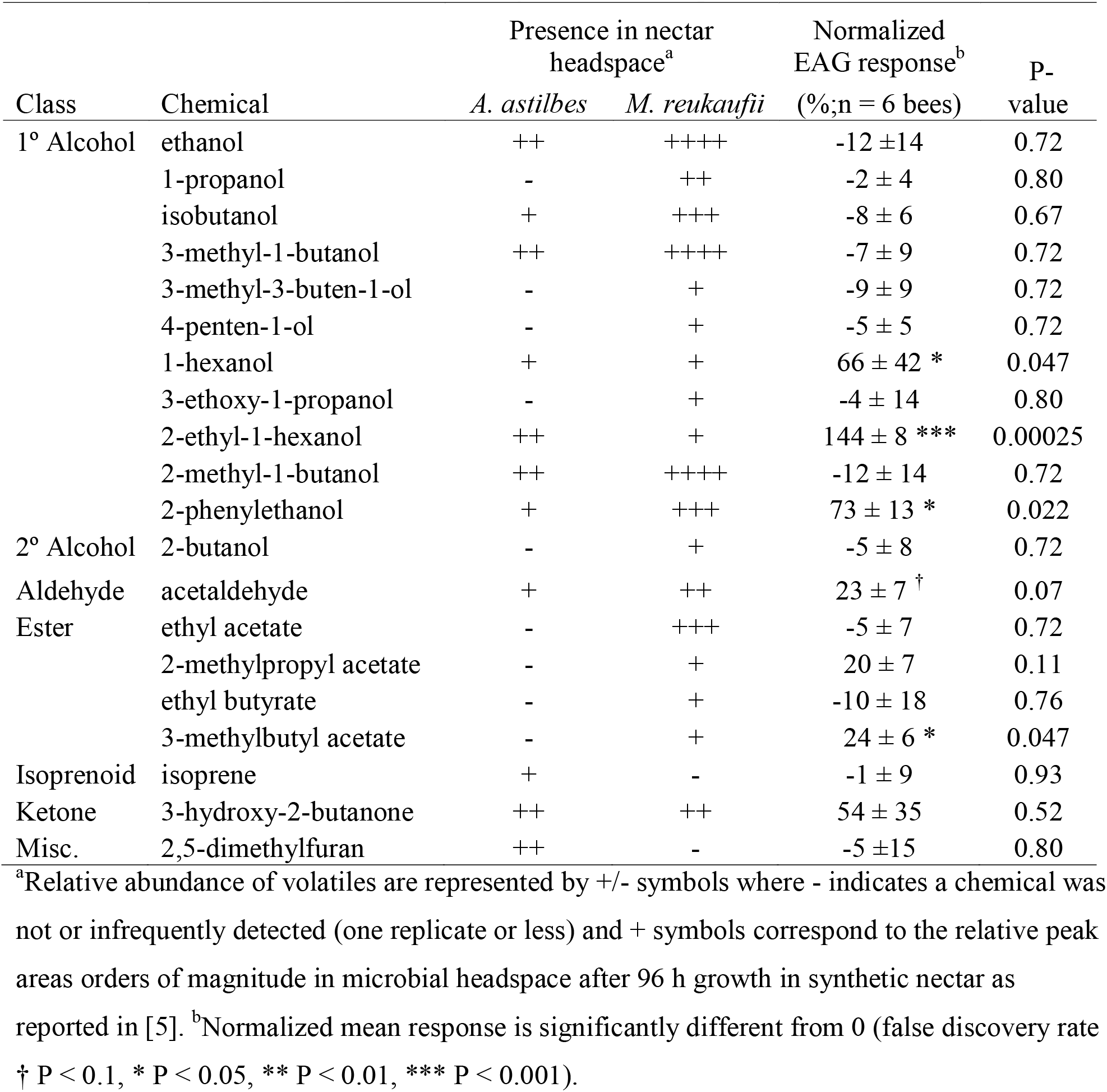
Volatile organic compounds produced by nectar-inhabiting microorganisms and their respective normalized mean bumble bee electroantennogram (EAG) response ± standard error (n=6) and corresponding false discovery rate corrected p-values.

| Class | Chemical | Presence in nectar headspace<sup>a</sup> |  | Normalized EAG response<sup>b</sup> (%; n = 6 bees) | P-value |
| --- | --- | --- | --- | --- | --- |
|  |  | *A. astilbes* | *M. reukaufii* |  |  |
| 1° Alcohol | ethanol | ++ | ++++ | -12 ± 14 | 0.72 |
| | 1-propanol | - | ++ | -2 ± 4 | 0.80 |
| | isobutanol | + | +++ | -8 ± 6 | 0.67 |
| | 3-methyl-1-butanol | ++ | ++++ | -7 ± 9 | 0.72 |
| | 3-methyl-3-buten-1-ol | - | + | -9 ± 9 | 0.72 |
| | 4-penten-1-ol | - | + | -5 ± 5 | 0.72 |
| | 1-hexanol | + | + | 66 ± 42 * | 0.047 |
| | 3-ethoxy-1-propanol | - | + | -4 ± 14 | 0.80 |
| | 2-ethyl-1-hexanol | ++ | + | 144 ± 8 *** | 0.00025 |
| | 2-methyl-1-butanol | ++ | ++++ | -12 ± 14 | 0.72 |
| | 2-phenylethanol | + | +++ | 73 ± 13 * | 0.022 |
| 2° Alcohol | 2-butanol | - | + | -5 ± 8 | 0.72 |
| Aldehyde | acetaldehyde | + | ++ | 23 ± 7 † | 0.07 |
| Ester | ethyl acetate | - | +++ | -5 ± 7 | 0.72 |
| | 2-methylpropyl acetate | - | + | 20 ± 7 | 0.11 |
| | ethyl butyrate | - | + | -10 ± 18 | 0.76 |
| | 3-methylbutyl acetate | - | + | 24 ± 6 * | 0.047 |
| Isoprenoid | isoprene | + | - | -1 ± 9 | 0.93 |
| Ketone | 3-hydroxy-2-butanone | ++ | ++ | 54 ± 35 | 0.52 |
| Misc. | 2,5-dimethylfuran | ++ | - | -5 ± 15 | 0.80 |

<sup>a</sup>Relative abundance of volatiles are represented by +/- symbols where - indicates a chemical was not or infrequently detected (one replicate or less) and + symbols correspond to the relative peak areas orders of magnitude in microbial headspace after 96 h growth in synthetic nectar as reported in [5]. <sup>b</sup>Normalized mean response is significantly different from 0 (false discovery rate † P < 0.1, \* P < 0.05, \*\* P < 0.01, \*\*\* P < 0.001).

### Experiment 2: Olfactory response of bumble bees to nectar-inhabiting microbes

To assess whether bumble bees exhibit innate preferences when exposed to volatile microbial metabolites, we used an olfactometer assay (Y-tube; Fig. S1) under red light. Naïve bumble bees housed at the University of California Davis were starved for 6 hours, then released individually into the Y-tube. For each bee, initial choice and the time spent in each arm was recorded, and the assay was repeated twice for each bee, with treatment direction reversed. These bees were both fed and treated similarly to those used for the EAG assays and a total of 32 bees were tested in this assay from two source colonies. For details, see electronic supplementary material S1 Methods.

### Experiment 3: Gustatory responses of bumble bees to nectar-inhabiting microbes

To assess gustatory responses of bumble bees (n=42 bees from two colonies) to nectar colonized by microbial taxa, we used both no-choice and choice feeding assays. For the no-choice assay, bees were housed in individual vials with modified lids that accommodated a feeding apparatus (Fig. S2). Vials were filled with 1 mL of either *Asaia*- or *Metschnikowia-* treated nectar, weighed, and bees were allowed to feed for 24 hr, after which tubes were reweighed to determine consumption. For details, see electronic supplementary material S1 Methods.

### Experiment 4: Effects of volatile and gustatory microbial cues on associative learning

Because bees exhibited marked differences in response to volatile and gustatory microbial cues (see Results below), we also assessed how exposure to gustatory cues influenced bee preference for volatiles (n=24 bees from two colonies). Individual foragers were subjected to the olfactometer assay (above), then a gustatory choice assay where individual bees were housed in a feeding chamber, consisting of ~9 cm of perforated tubing, with feeding vials on either end of the chamber (Fig. S3) for 24 hr. Vials were weighed to determine nectar consumption. Bees were then subjected to a second olfactometer assay.

### Statistical analyses

To assess which compounds were detected by bumble bees (*Experiment 1*), we used *t*-tests with false discovery rate correction to examine if normalized EAG responses were significantly different from zero (i.e., no detectable response). To determine if bee preference differed between microbes, data from *Experiment 2* were analyzed using a binomial test for first choice. A linear mixed-effect (LME) model [22] was used for time spent in each arm, with microbial treatment as a fixed effect, and bee individual as a random effect. For gustatory cues (*Experiment 3*), we used a t-test to assess how nectar consumption was affected by the nectar treatment. For *Experiment 4*, we fit a LME model with proportion of time spent in olfactometer arms as the response variable, nectar treatment, choice test order, and their interaction as fixed effects, and bee individual as a random effect. Bumble bee feeding responses were also analyzed with a LME model, with amount consumed as the response variable, nectar treatment as a fixed effect, and bee individual as a random effect. All analyses were performed in R (v. 3.5.2) [23].

## Results

Bumble bee antennae responded to a subset (4/20) of volatile metabolites tested through EAG (*Experiment 1*; Table 1) at 0.4 μmol, including 1-hexanol, 2-ethyl-1-hexanol, 2-phenylethanol, and 3-methylbutyl acetate (i.e., isoamyl acetate, isopentyl acetate). The alcohol 2-ethyl-1-hexanol elicited the strongest EAG depolarization response, surpassing that of the positive control (0.4 μmol geraniol).

Volatile blends emitted by nectar-inhabiting microbes also influenced bee behavior. Naïve bees on average spent ~two-thirds of their time in Y-tube arms containing *Asaia*-produced volatiles (Figure 1A; *F_1,64_*=21.52, *P*<0.0001), although no difference was found for first choice (*P*=0.67). In contrast, bees consumed approximately 50% less Asaia-conditioned nectar than *Metschnikowia* nectar (Figure 1B; *t_29.5_*=-2.70, *P*=0.011) in a no-choice assay (*Experiment 3*).

**Figure 1.**
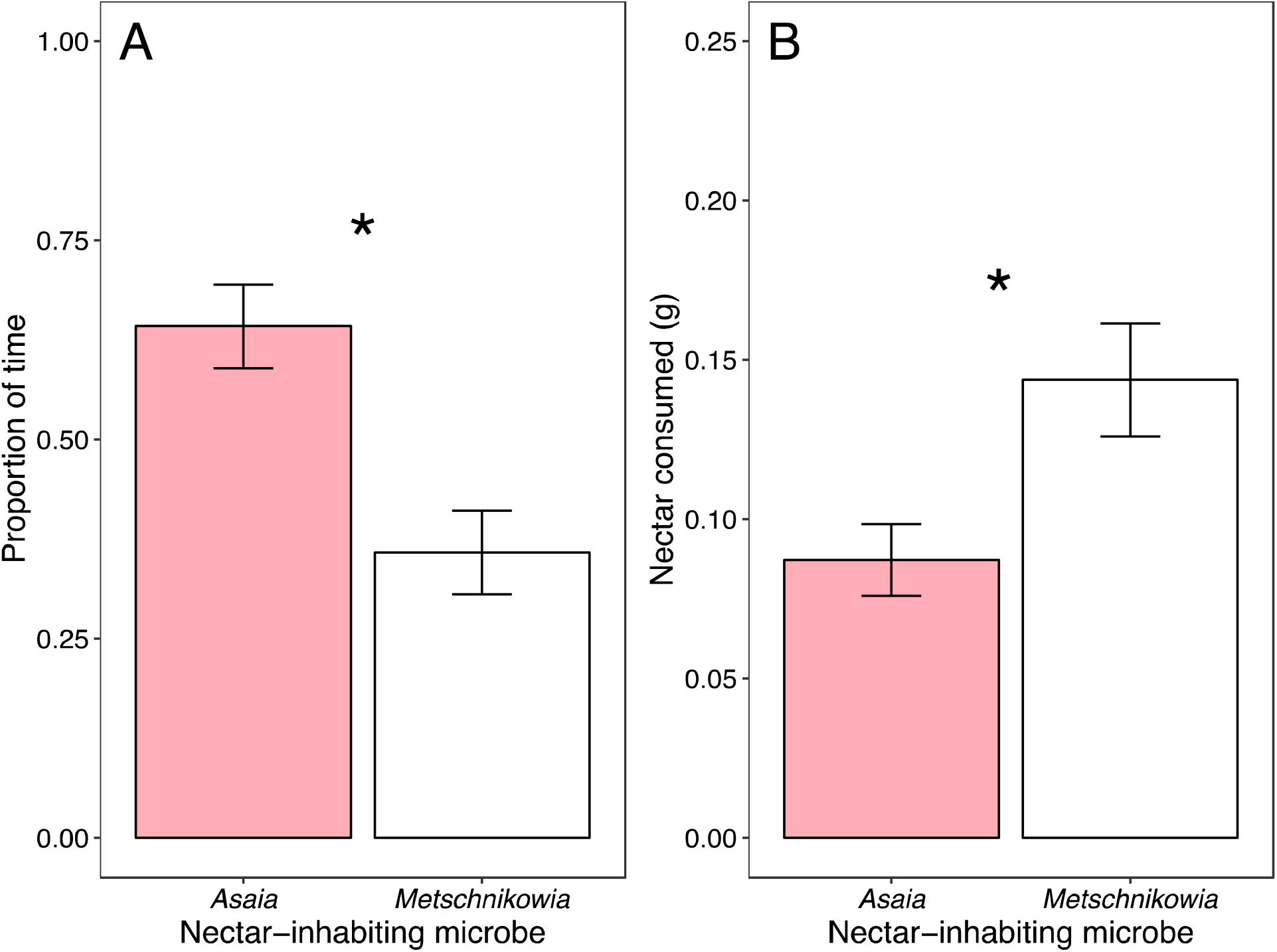
Behavioral (A) and gustatory (B) responses of bumble bees to artificial nectar colonized by nectar-inhabiting microbes and the volatile organic compounds they emit.

Mirroring earlier results, bumble bees across both choice tests performed in *Experiment 4* spent ~15% more of their time in the Y-tube arm assigned to *Asaia* compared to that of *Metschnikowia* (*F_1,163_*=9.09, *P*=0.003). During the feeding assay, bees again consumed on average nearly double the amount of *Metschnikowia-conditioned* nectar compared to *Asaia* (*F_1,46_*=12.29, *P*=0.001). After experiencing gustatory cues in the feeding assay, bees reduced the frequency with which they chose the *Asaia* volatile blend, increasing both the proportion of ‘no choice’ and that for the yeast arm, as well as the amount of time spent (albeit not significant) in the *Metschnikowia* arm of the olfactometer.

## Discussion

Microbes commonly inhabit food resources, contributing both volatile and non-volatile metabolites that can function to inform foragers as to their location and quality. We found distinct effects of olfactory vs. gustatory cues produced by two common, nectar-inhabiting microbes on bumble bee behavior and nectar consumption. The difference in bee response to olfactory vs. gustatory cues suggests that these cues are not reinforcing, which may complicate pollinator foraging and learning based on microbial metabolites [24]. Further, bee preference for *Asaia* volatiles decreased after exposure to gustatory cues and feeding, suggesting behavior modification. We suspect that acetic acid produced by *Asaia* (but not *Metschnikowia*), although not detectable in our volatile screening, may be aversive to bees. In natural systems, bees likely develop associations between microbial chemosensory cues through repeated exposure to the scent and taste of yeast or bacterial-colonized nectar. However, our findings, and recent experimental results [24], suggest that microbial signals may be more difficult to learn than other sensory combinations. Such difficulties may manifest to affect learned preferences, floral constancy and the quantity and quality of benefits exchanged in these mutualistic interactions.

Collectively, our results indicate that volatile and non-volatile microbial metabolites have significant potential to shape interspecific, plant-pollinator signaling. In remains to be determined whether pollinators benefit from microbial-derived cues can translate to improved foraging efficiency, or whether such cues may be more exploitative, and benefit microbes that rely upon pollinator dispersal to reach new floral habitats [25]. Such outcomes may hinge on both the identity and density of the microbial species encountered, where varied immigration histories can give rise to divergent microbial communities both within flowers of a host and among other species. Our results demonstrate that future investigations on the evolutionary ecology of floral signaling should consider multiple ways in which microbes influence host phenotype and the innate and learned response of pollinators.

## Authors’ contributions

R.S., C.R., J.B., and R.V. conceived the study. R.S., C.R., and I.M. collected data, while R.S. and C.R. performed statistical analyses and drafted the manuscript. All authors contributed to manuscript editing, gave final approval for publication, and agree to be held accountable for the worked performed therein.

## Acknowledgements

We thank M. Handy, A. Khan, I. Munkres, H. Pathak, and M. Anderson for laboratory assistance, and B. Forguson for greenhouse assistance.

