## Supplementary material for "Microbial metabolites mediate bumble bee attraction and feeding"

**S1 Methods**

***Experiment 1: Electroantennographic bioassay***

Bumble bees (*n* = 6 bees tested for each metabolite) were collected each day from a commercial colony housed in a greenhouse located at the USDA‐ARS Center for Medical, Agricultural and Veterinary Entomology (Gainesville, FL, USA). Before experiments, bees were fed a pollen and sugar water paste (1:1 pollen and 30% sucrose) *ad libitum*. Immediately before bioassays, individuals were captured in a trimmed 15 mL centrifuge tube and secured from behind with cotton. Under a low power stereo‐microscope, both antennae were excised at the scape using micro‐scissors and mounted with the same orientation on a forked probe (Syntech, Kirchzarten, Germany; internal gain 10×) with electrode gel (Parker Laboratories, Fairfield, NJ, USA). The prepared probe was mounted in the humidified constant air and volatile sample tube and allowed to equilibrate to the air flow for 2–3 min (until signal stabilized). Sample puffs were delivered at 1‐min intervals to allow antennae to re‐equilibrate after exposure. Antennal responses, which indicate detection rather than attraction or repulsion, were recorded with the Autospike software (Syntech, v.3.9). Maximum antennal responses were selected within a 2 sec window after sample puff and recorded as absolute values.

Odor samples were prepared using 6 mm grade AA discs (Whatman, Maidstone, United Kingdom), loaded with a 0.4 μmol solution containing an individual metabolite. The selected concentration was informed by consistent bumble bee antennal responses to known social bee pheromones at this dose across individuals screened in preliminary tests. The carrier solvent (pentane or acetone, depending on metabolite solubility) was allowed to evaporate for 2 min and then the filter paper was placed within a Pasteur pipette trimmed at the tapered end to a final length of 6.5 cm and sealed with Parafilm (Bemis NA, Neenah, WI, USA). Odors were directed through the electroantennographic apparatus containing the mounted antennae and probe using both a 0.5 sec pulse flow (300 ml min^−1^) and a humidified continuous flow (125 ml min^−1^). Ambient electrical interference was eliminated via use of a Faraday cage. To account for variability in response among individuals and over time as antennae dehydrate, samples were bracketed by blanks (20 μL carrier solvent corresponding to sample and bioassay disc) and standard stimuli (0.4 μmol geraniol dissolved in corresponding carrier solvent and bioassay disc). For each sample puff, mean blank responses were subtracted from sample and standard stimuli responses and the sample antennal response values were then normalized to the mean blank-corrected standard stimuli response, defined as 100% as follows:

$$\frac{\left[ \left| S \right|-\left| B \right| \right]}{\left[ \left| P \right|-\left| B \right| \right]} \times100\%=\% response$$

Where *S* (µV) is the maximum recorded response from a given sample puff, and *B* (µV) and *P* (µV) are the mean response of the preceding and following blanks and standard stimuli, respectively.

***Experiment 2: Olfactory response of bumble bees to nectar-inhabiting microbes***

Prior to each test, individual bumble bees were housed in ventilated, plastic vials in the dark without food for 6 hr. Each individual was then released in the entrance arm of the Y-tube. Test sessions were carried out in the dark under red light, to both favor bumble bees' use of olfactory cues and allow for observation of bee behavior in real time. During the test, two olfactory stimuli were delivered, each coming from one of the two arms of the Y-tube. Stimuli (i.e. microbial-conditioned nectar solutions) were deposited on 1-cm^2^ pieces of filter paper (Whatman) and then inserted into vacuum trap bubblers. Carrier tubes extending from these bubblers were connected to each arm of the Y-tube. Moistened, pure air, sourced from a medical-grade cylinder, was pulsed at 20 mL min^−1^ from the dead end of each arm through these carrier tubes, allowing the olfactory stimuli to flow towards the decision area, defined as the area where the bumble bee had to make a choice. Between tests, the Y-tube was carefully cleaned using an ethanol solution and allowed to dry before subsequent use.

For each bee tested, we recorded its first choice, as well as the amount of time spent in each arm of the Y-tube during 5 min of observation. Each test was performed twice for each bee, with the arm assigned for each olfactory stimuli flipped in the subsequent assay to control for any positional effects. A total of 32 bees were tested in this assay from two source colonies.

***Experiment 3: Gustatory responses of bumble bees to nectar-inhabiting microbes***

To assess gustatory responses of bumble bees to nectar colonized by microbial taxa, we used both no-choice and choice feeding assays. For the no-choice assay, bees were housed in individual vials (26 mm x 51 mm; Fisher Scientific) with modified lids that accommodated a feeding apparatus (Fig. S2). This apparatus consisted of a 1.7 mL microcentrifuge tube with lid removed, and ~0.14 g of cotton wick. At the start of the assay, feeding vials were filled with 1 mL of either *Asaia-* or *Metschnikowia-*treated nectar, and bees were allowed to feed for 24 hr. At the end of the assay, each tube was weighed to determine the amount of treated nectar consumed. A subset of cold-anesthetized bees were also weighed after the 6 hr starvation period to account for any differences in feeding that could be attributed to bee size, however no significant effect was detected and this factor wasn’t included in further analyses (see below). A total of 42 bees were tested in this assay from two source colonies.

**S1 Figures**

. . .

Olfactometer (Y-tube)

- Main channel and arms: 1.9 cm H

- Main channel: 9 cm L

- Arms: 3.8 cm L

3.8 cm of 0.95 cm ID 180 PVC autoclavable tubing

mesh, taped to end of tubing

vacuum trap bubbler; filter paper with 10 μL of treatment goes in bottom of bubbler

5 cm of 0.95 cm ID 180 PVC autoclavable tubing

0.95 cm ID 180 PVC autoclavable tubing connected to a pressure gauge and zero air tank

12 cm glass y-joint connector

**Figure S1.** Schematic drawing of Y-tube olfactometer. Filter paper, treated with respective microbial-condition nectar solutions were placed inside vacuum trap bubblers. Bumble bees entered the main channel of the olfactometer, and first choice and time spent in either arm of the Y-tube was recorded under red light.

1.5 mL flat-top centrifuge tube, cap removed

0.14 g cotton ball/wick

1 mL treatment

snap-on lid with breathing holes and an “x” cut into it, to easily insert and remove centrifuge tube

26 mm x 51 mm vial

**Figure S2.** Schematic of feeding vial used in no-choice feeding assay to assess bumble bee feeding responses to microbial gustatory cues.

8.9 cm L x 3.8 cm ID

180 PVC tubing

1.5 mL centrifuge tube, lid removed

parafilm

0.14 g cotton ball/wick

1 mL treatment

**Figure S3.** Schematic of feeding tube used in choice assay to assess bumble bee responses to microbial gustatory cues.
